# Gut microbiota and faecal cytokine profiles of rural Cambodian infants and links to early life nutrition and diarrhoeal episodes

**DOI:** 10.1101/2023.07.24.550269

**Authors:** Matthew J Dalby, Raymond Kiu, Iliana R Serghiou, Asuka Miyazaki, Holly Acford-Palmer, Rathavy Tung, Shabhonam Caim, Sarah Phillips, Magdalena Kujawska, Mitsuaki Matsui, Azusa Iwamoto, Bunsreng Taking, Sharon E Cox, Lindsay J Hall

## Abstract

The gut microbiota of infants in low-middle income countries like Cambodia remain underrepresented in microbiome research. This study aimed to explore the faecal gut microbiota composition and faecal cytokine profiles in a cohort of infants living in a rural province of Cambodia and explore the impact of sample storage conditions and infant environment on microbiota composition. Faecal samples collected at three time points from 32 infants (96 samples in total) after 7 months of age were analysed using 16S rRNA amplicon sequencing to determine the composition of the microbiota. Bacterial strains were isolated and subjected to whole genome sequencing and genomic analysis and concentrations of faecal cytokines were also measured. Initially, we compared the effects of two sample collection methods due to the challenges of faecal sample collection and storage in a rural location. Storage of faecal samples in a DNA preservation solution retained a greater abundance of *Bacteroides*. Analyses of microbiota composition of samples stored in DNA preservation solution indicated that *Bifidobacterium* was the most abundant genus with *Bifidobacterium longum* the most abundant species, particularly in breastfed infants. Most infants had detectable pathogenic taxa indicating frequent pathogen exposure, with *Shigella* and *Klebsiella* more abundant in infants with recent diarrheal illness. We did not detect antibiotic-associated perturbations in the gut microbiota, and no associations were found between the gut microbiota and infant growth. Genomic analysis of isolated strains indicated the presence of gene clusters encoding the ability to digest human milk oligosaccharides in *B. longum* and *Bifidobacterium breve* isolates. The presence of antibiotic-resistant genes was also identified in potentially pathogenic species, as well as in beneficial genera including *Bifidobacterium*. Faecal cytokine analysis showed higher concentrations of Interlukin-1alpha and vascular endothelial growth factor in breastfed infants, which may influence the infant gut mucosal immune system. This study provides insights into an underrepresented population of rural Cambodian infants, emphasising the impact of pathogen exposure and breastfeeding on gut microbiota composition and faecal immune profiles.

## Introduction

Research on the gut microbiota has primarily focussed on populations in high income countries (e.g. Europe and North America) (Abdill et al., 2022), and the problems and limitations this bias is introducing to microbiome research is gaining increasing attention (Browne et al., 2024). Although more recent studies have increased representation of populations in low- and low-middle income countries (LMICs), infant studies lag behind those working on adult cohorts (Porras and Brito 2019). This is important for defining what is a ‘healthy’ human gut microbiota (Lackey et al., 2019) across individuals in LMICs (Porras and Brito, 2019), and because optimal infant development is particularly important for infants from LMICs due to increased risk of (mal)nutrition and infection. While a small proportion of microbiota samples studied globally so far have come from South Asia, including India and Bangladesh (Abdill et al., 2022), research into the gut microbiota of people living outside these countries remains very limited. Moreover, due to the logistics of sample collection and storage in rural areas, research in LMICs has often focused on individuals in urban areas, neglecting key infant populations. Cambodia in particular has a scarcity of microbiota studies and given that this country has such a high prevalence of infants classified as stunted/wasted, and has amongst the highest infant mortality rates in Southeast Asia, with infections accounting for many of these deaths (Cambodia Demographic and Health Survey, 2022), new studies targeting Cambodian infant populations are needed. Previous research focusing on the gut-associated microbiota of people in Cambodia has been limited with studies focusing on particular bacterial pathogens, such as antimicrobial resistance in *Enterobacterales* in faecal samples from babies and children (Auguet et al., 2021), the epidemiology of antibiotic resistance in *Escherichia coli* and *Klebsiella pneumoniae* in children (van Aartsen et al., 2019), and effects of iron supplementation in Cambodian women (Finlayson-Trick et al., 2023). However, analysis of the composition of the gut microbiota as a whole in infants from Cambodia has not previously been published.

While the nutritional status of infants is improving in Cambodia, children under the age of 5 in rural areas still face challenges, with 22% suffering linear growth faltering or ‘stunting’ (Cambodia Demographic and Health Survey, 2022). This is also true for infants (<1 year) in rural Cambodia who also suffer from similar levels of growth stunting at 19.2% (Miyazaki et al., 2021), with a high proportion also lacking access to appropriate hygiene facilities. This is coupled with a high proportion reportedly experiencing symptoms of diarrhoea, fever or cough and frequent use of unregulated antibiotic treatment to treat these childhood illnesses (Miyazaki et al., 2020). Malnutrition has previously been linked to differences in the gut microbiota of children from several different low income countries (Robertson, 2020; Surono et al., 2021), although research into these associations is lacking for children from Cambodia.

The establishment of the gut microbiota during infancy is important aspect of healthy infant development with roles in immune, metabolic, endocrine, and other host developmental pathways (Robertson et al., 2019). Vaginally-delivered, breast fed infants are characterised by their very high abundance of the bacterial genus *Bifidobacterium.* This keystone microbiota member orchestrates wider microbiome structuring and modulates host physiology through the breakdown of complex dietary components such as human milk oligosaccharides (HMOs) in breast milk and plant-based carbohydrates, leading to production of key metabolites that facilitate immune system development and maturation and cognitive responses (Henrick et al., 2021). Previous research indicates that LMIC infant microbiomes are characterised by *Bifidobacterium* species and strains, although these are often different genetic variants when compared to those infants from high income country settings (Lackey et al., 2019; Taft et al., 2022).

Infants born into resource-poor environments can have difficulties in establishing a healthy gut microbial ecosystem (Lang & MAL-ED, 2018) due to multiple factors such as birth and breast feeding practices, antibiotic exposure, maternal microbiota, diet, socio-economic status, host genetics, agricultural dependence, urbanisation, access to clean water and hygiene practices (Porras and Brito, 2019; Robertson et al., 2019). Infant microbiomes of those in LMICs are reportedly higher in *Segatella* (previously *Prevotella*) species, associated with both positive and negative health outcomes, and may correlate with higher plant based, fibre rich diets (Porras and Brito 2019). The phylum Pseudomonadota (previously Proteobacteria) that includes a variety of pathogenic and opportunistic pathogens, is highly prevalent in microbiomes of those in LMICs (Porras and Brito 2019). These factors may negatively affect the composition of the early infant gut microbiota (Robertson et al., 2019), and may drive these observed microbiome differences across populations (Porras and Brito 2019).

In this study we carried out an exploratory analysis of the gut microbiota in faecal samples collected from 32 infants that are part of the Nutrition for Health of Aka-chan (baby in Japanese) and Mamas’ cohort (NHAM) based in Cambodia, which included background data on infant diet, health, and living conditions. This provides an opportunity to explore the composition of the gut microbiota in rural Cambodian infants that has not previously been investigated. The samples from this cohort are unusual due to the rural location in which these infants live, with poor access to hygiene facilities, high prevalence of pathogen exposure, and frequent antibiotic usage. We analysed the effects of faecal sample storage on the composition of the microbiota of samples collected under suboptimal conditions. Then described the infant faecal microbiota composition and explored its associations with environmental factors such as infant diet and growth. Individual faecal bacteria cultured and isolated and their genomes sequenced to identify the presence of antibiotic resistance and HMO genes. Finally, the concentrations of faecal cytokines were measured to identify any associations with the gut microbiota. We aimed to explore the composition of gut microbiota of these infants and its associations with their environment.

## Results

The longitudinal infant faecal samples analysed in this study were sourced from 32 infants representing a subset of infants that are part of the NHAM birth cohort based in the rural Kampong Cham province in southeast Cambodia, and approximately 60 kilometres from the provincial capital Kampong Cham (Miyazaki et al., 2020, 2021) (**Table 1**). Samples were collected from infants living in a riverside community stretching along approximately 15.5 km of riverbank of the Mekong River from Khbop Ta Nguon to Peam Koh Sna (Miyazaki et al., 2021). The infants were living in a rural area, with most families being small-scale farmers.

**Table 1:** Infant baseline characteristics.

| Variable | Infants |
| --- | --- |
| n | 32 |
| Age (days) (mean (SD)) | 214.9 (37.3) |
| sex = Female (%) | 15 (46.9) |
| Breast feeding = No (%) | 9 (28.1) |
| Complementary feeding = No (%) | 17 (53.1) |
| Feeding bottle use = Yes (%) | 21 (65.6) |
| Low birth weight (<2.5kg) (%) |  |
| Normal | 29 (90.6) |
| Unknown | 2 ( 6.2) |
| Low | 1 ( 3.1) |
| Growth = Stunted (%) | 3 ( 9.4) |
| Length-for-age z-score (mean (SD)) | -0.9 (0.8) |
| Underweight z-score (mean (SD)) | -0.6 (0.8) |
| Weight-for-length (mean (SD)) | 0.0 (1.0) |
| Hand washing score (mean (SD)) | 5.2 (1.6) |
| Improved toilet facility = Yes (%) | 22 (68.8) |
| Household toilet facility = Septic tank (%) | 22 (68.8) |
| Number in household (mean (SD)) | 5.8 (2.2) |
| Antibiotic use (previous 7 days) = No (%) | 28 (87.5) |
| Any recent illness (previous 7 days) = No (%) | 11 (34.4) |
| Diarrheal illness (previous 7 days) = No (%) | 26 (81.2) |
| Fever (previous 7 days) = No (%) | 17 (53.1) |
| Cough (previous 7 days) = No (%) | 20 (62.5) |

### Sample storage conditions influence gut microbiota profiles

Sample collection and storage is challenging in areas without the facilities to immediately freeze faecal samples after collection. To compare the preservation of DNA in samples under different conditions, half of each faecal sample was immediately placed into a DNA preservation solution (DNAShield), with the remaining half remaining untreated. The faecal samples were collected in a rural location with approximately 2 days between the sample being produced and final laboratory storage. The untreated half of each sample was transported in iceboxes and then initially frozen at - 20°C and then later stored at -80°C. The half of each sample stored in DNAShield were transported at ambient temperature and then stored refrigerated at 4°C. Three, biweekly samples from 32 infants were collected (total of 96 samples), with half split between each storage condition. Both frozen and DNAShield paired infant gut microbiota samples underwent 16S rRNA amplicon sequencing to analyse their bacterial composition.

Out of the 96 paired samples, a total of 94 were successfully sequenced when stored in DNAShield, while only 70 were successfully sequenced from Frozen samples (**Supplementary File 1**). There were 69 pairs of samples with successful 16S rRNA amplicon sequencing from both the Frozen and DNAShield portion of the same faecal sample. The resulting bacterial genus composition was compared between these 69 pairs of samples (**Figure 1**). While the total number of bacterial genera detected was similar between frozen and DNAShield samples (**Figure 1A** ), microbiota diversity calculated using the Shannon index and Inverse Simpson index was higher in DNAShield samples than paired frozen samples (**Figure 1B-C** ). *Bacteroides*, *Escherichia, Parabacteroides, and Lachnoclostridium* were the four bacterial genera with the largest median difference in relative abundance between storage conditions that were higher in DNAShield samples compared to frozen samples (**Figure 1D-G; Supplementary File 1** ). *Bacteroides* showed the largest difference in favour of DNAShield storage with a median relative abundance of 0.4% in frozen samples and 8.6% in DNAShield samples (**Figure 1D; Supplementary File 1** ). The relative abundance of *Bifidobacterium*, *Streptococcus*, *Enterococcus,* and *Actinomyces* were the genera with the largest difference and significantly higher abundance in favour of frozen samples (**Figure 1H-K; Supplementary File 1** ). *Bifidobacterium* comprised a median relative abundance of 63% in frozen samples and 46% in DNAShield samples (**Figure 1H; Supplementary File 1** ). Further analysis of bacterial microbiota composition in this study was performed using samples stored in DNAShield due to the larger number of samples with successful sequencing results and the higher abundance of individual bacterial genera surviving in DNAShield stored samples.

**Figure 1.**
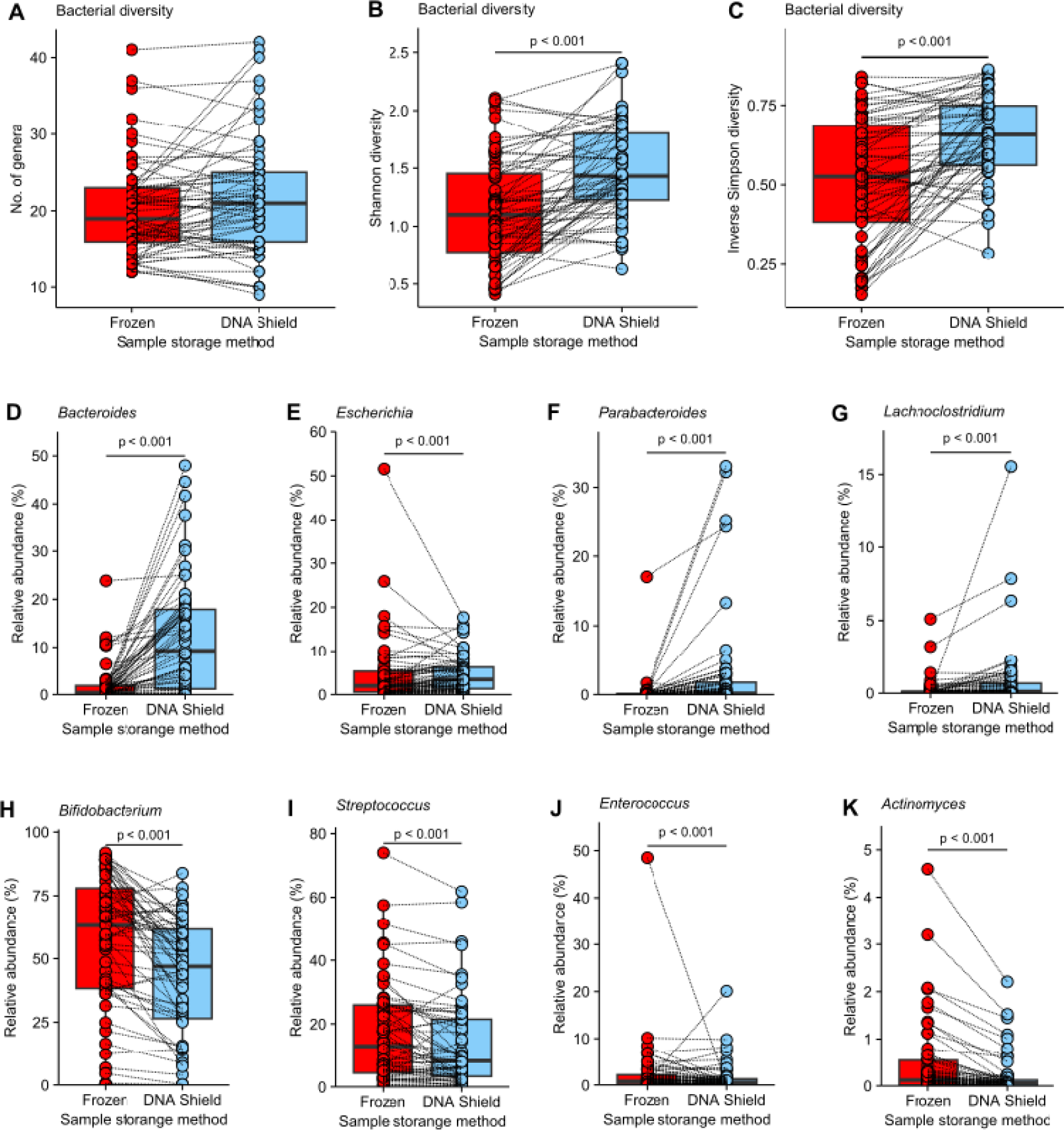
Differences in genus abundance between paired frozen samples and DNAShield stored samples. (A) The number of genera detected in each sample; (B) The Shannon diversity index of each sample; (C) The Inverse Simpson diversity index of each sample; Relative abundance between paired sampled after different storage conditions of (D) *Bacteroides*; (E) *Escherichia*; (F) *Parabacteroides* ; (G) *Lachnoclostridium*; (H) *Bifidobacterium*; (I) *Streptococcus*; (J) *Enterococcus*; (K) *Actinomyces*. Box plots show median and interquartile range with circles showing individual samples. Dotted lines link together paired Frozen and DNAShield samples from the same infant sample. Statistical significance between groups was tested using a Wilcoxon signed rank test with correction for multiple testing amongst bacteria genera using the false discovery rate method.

These results indicate that sample storage conditions selectively can alter the abundance of different genera in analysis of bacterial composition with significantly higher abundance of *Bacteroides* and *Parabacteroides* remaining when samples were stored in DNA preservation solution. This shows that DNA preservation solution can preserve genus abundance in final composition analysis even when samples are initially transported at ambient temperatures and then stored refrigerated and not frozen.

### Bacterial genus composition of the infant gut microbiota is influenced by breastfeeding and diarrheal illness

The gut microbiota of infants was dominated by *Bifidobacterium,* which comprised a median relative abundance of 48.7% (**Figure 2A** ). In addition, other genera including *Bacteroides, Blautia, Erysipelatoclostridium, Escherichia, Lactobacillus, Megamonas, Parabacteroides, and Streptococcus*, comprised the top ten most abundant genera, and formed the majority of the gut microbiota in these infants (**Figure 2A**).

**Figure 2.**
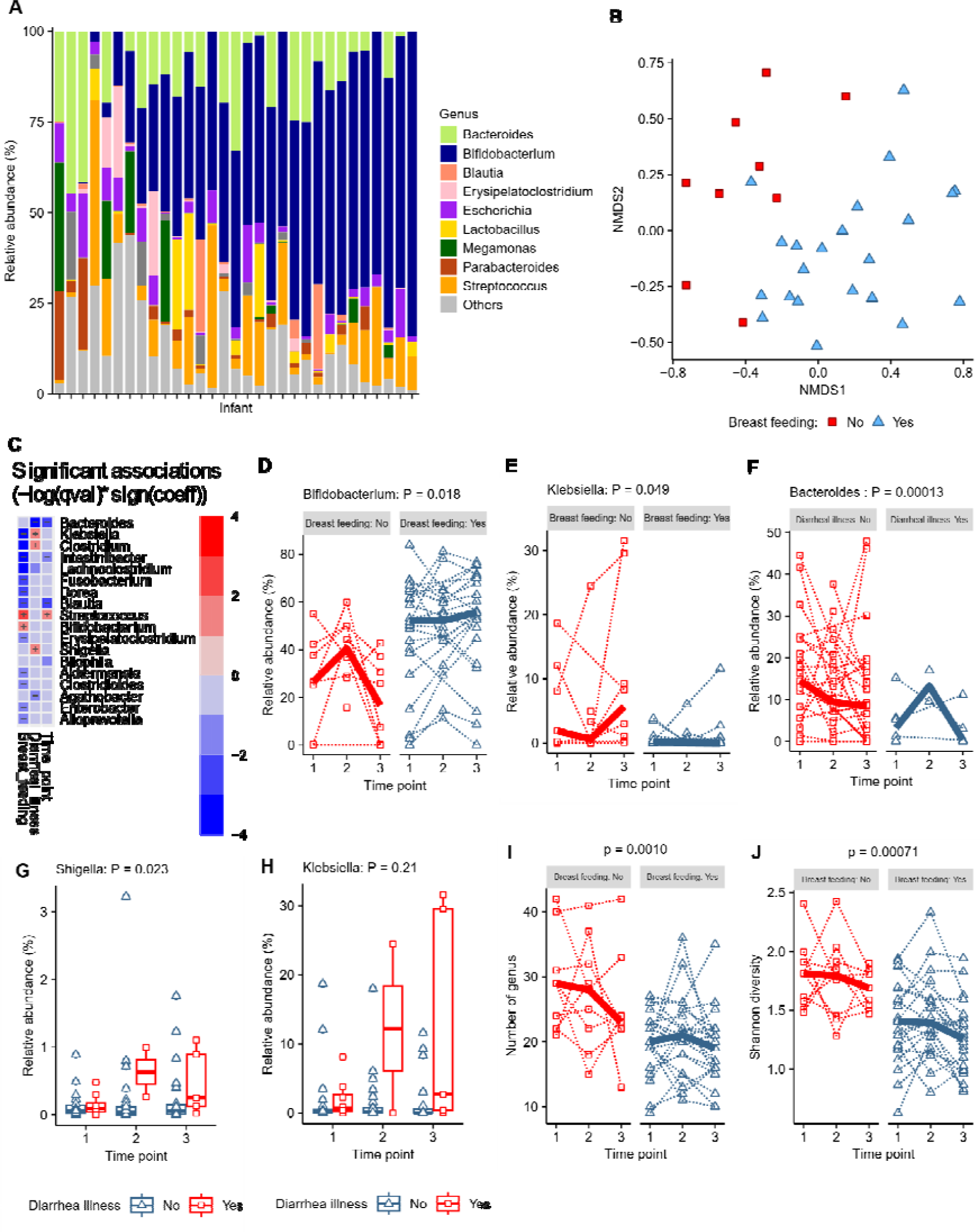
Genus composition and associations of genus composition with breast feeding status and diarrheal illness. (A) The ten most abundant bacterial genera across all samples ordered on the x-axis by *Bifidobacterium* abundance in each infant; (B) Non-metric multidimensional scaling (NMDS) plot representing beta-diversity of infant genus microbiota at time point 1 clustered by similarity and coloured by breastfeeding status with arrows showing top ten bacterial genus diving separation of samples; (C) Heat map showing significant associations determined by MaAsLin2 analysis between genus and breast feeding, diarrheal illness and time point; (D) The relative abundance of the genus *Bifidobacterium* by breastfeeding status; (E) The relative abundance of the genus *Klebsiella* by breastfeeding status; (F) The relative abundance of the genus *Bacteroides* by breastfeeding status; (G) The relative abundance of the genus *Shigella* by diarrheal illness within the previous 7 days; (H) The relative abundance of the genus *Klebsiella* by diarrheal illness within the previous 7 days; (I) The number of genera by breast feeding status; (J) The Shannon diversity index by breast feeding status; Points on plots show individual infant data and dotted lines on spaghetti plots link individual infants across time points with the bold line showing the group median.

An exploratory analysis was carried out using metadata available for the infants from which samples were collected. This included cross-sectional data collected at the start of the study before collection of the faecal sample at time point 1 and additional information collected at each sample collection visit (**Supplementary File 2** ). The exploratory analyses used non-metric multidimensional scaling (NMDS) and sample beta-diversity to cluster the samples by the similarity of their overall microbiota composition at time point 1. Associations between the infant metadata with the beta-diversity of the gut microbiota was analysed using Envfit and significant associations illustrated on the NMDS plot (**Figure 2B** ). Only breast-feeding status was significantly associated with microbiota beta-diversity with clear separation in clusters between infants breast fed and those infants not breast fed (**Figure 2B**).

To identify associations between the relative abundance of individual genera and infant metadata an analysis using the MaAsLin2 package was carried out that included the samples from all three time points, with the time point variable included as a random effect (**Figure 2C; Supplementary File 3** ). When background metadata variables were analysed individually as a fixed effect, both breastfeeding status and recent diarrhoeal illness within the previous seven days showed significant associations with genus abundance, which was maintained when both variables were included together as a multivariable analysis (**Figure 2C** ). Genera with differential abundance that was significantly associated with breast-feeding and/or diarrheal illness are shown as a heat map with positive and negative symbols indicating the direction of association (**Figure 2C** ). The genera positively associated with breast-feeding were *Bifidobacterium* and *Streptococcus* (**Figure 2C**; **Figure 2D**), whilst those with the strongest negative associations were *Klebsiella* and *Clostridium* (**Figure 2C**; **Figure 2E** ). Breast-feeding status remained unchanged for individual infants across the three time points. At time point 1 the mean age of breast fed infants was 207 days and non-breast fed infants 214 days. Including age as a variable in the analysis did not alter the associations with breastfeeding. Other measures of infant diet, the use of a feeding bottle or complementary feeding, were not associated with genus abundance. MaAsLin2 analysis also showed that that recent diarrheal illness within the previous seven days was associated with a lower relative abundance of *Bacteroides* (**Figure 2F** ) and higher relative abundance of *Clostridium*, *Klebsiella* and *Shigella* (**Figure 2G-H** ). *Shigella* was detected in 78% of samples, *Klebsiella* in 82% and *Clostridium* in 65% indicating that these infants have a high prevalence of these potentially pathogenic genera in their gut microbiota (**Supplementary File 4**). Breast feeding was the only variable associated with differences in microbial diversity with the total number of bacterial genus (**Figure 3I** ) and the Shannon diversity lower in breast fed infants than non-breast fed infants (**Figure 3J**).

**Figure 3.**
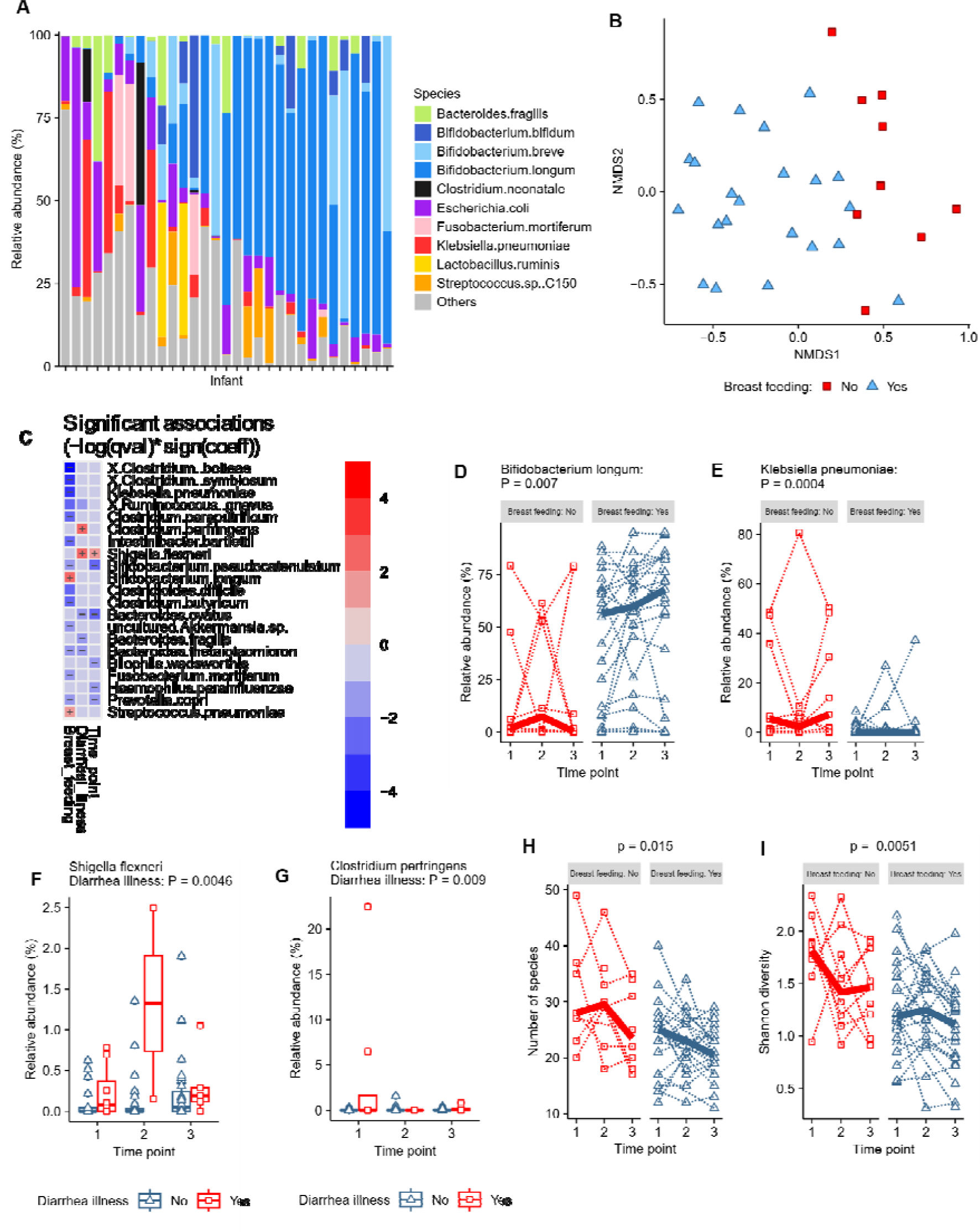
Species composition and association of species composition with breast feeding and diarrheal illness. (A) The top ten most abundant bacterial species across all samples at time point 1 ordered by total *Bifidobacterium* species abundance; (B) Non-metric multidimensional scaling (NMDS) plot representing beta-diversity of infant species microbiota at time point 1 clustered by similarity and coloured by breastfeeding status with arrows showing top five bacterial genus diving separation of samples; (C) Heat map showing significant associations determined by MaAsLin2 analysis between species and breast feeding, diarrheal illness and time point; (D) The relative abundance of the *Bifidobacterium longum* by breastfeeding status; (E) The relative abundance of the *Klebsiella pneumoniae* by breastfeeding status; (F) The relative abundance of the *Shigella flexneri* by diarrheal illness within the previous 7 days; (G) The relative abundance of *Clostridium perfringens* by diarrheal illness within the previous 7 days; (H) The number of species by breast feeding status; (I) The Shannon diversity index by breast feeding status, Points on plots show individual infant data and dotted lines on spaghetti plots link individual infants across time points with the bold line showing the group median.

Only three infants included in the analysis were classified as having stunted growth with a height- for-age z-score of less than 2 and this was not associated with microbiota composition. Tested as continuous measures, length-for-age z-score , underweight z-score, and weight-for-length z-score were also not associated with microbiota composition. However, this number of infants was too small for robust testing preventing any strong conclusions being drawn from this result. Measures of household hygiene facilities and total number of people in the household were not associated with microbiota composition. Of particular note was the lack of an association between genus composition and recent antibiotic use within the previous 7 days (**Supplementary File 3**).

These results indicate that breast feeding was the most influential factor shaping the gut microbiota composition in favour of increased *Bifidobacterium* abundance. These results also show that recent diarrheal illness was associated with higher relative abundance of the potentially pathogenic genera *Clostridium*, *Klebsiella* and *Shigella*.

### Bacterial species composition of the infant gut microbiota is influenced by breastfeeding status

The same analysis carried out on genus level composition was carried out on the data at the species level with the limitation that species classifications using 16S rRNA amplicon data are less robust than genus (**Supplementary File 5** ). The gut microbiota at the species level was dominated by a single *Bifidobacterium* species classified as *Bifidobacterium longum* (**Figure 3A** ). Other abundant species detected were members of those genera identified in (**Figure 2A**) with two other species of *Bifidobacterium* identified as *B. breve* and *B. bifidum*, while *Escherichia coli*, *Klebsiella pneumoniae*, *Bacteroides fragilis* and an unclassified *Streptococcus* comprised the other abundant species present.

Repeating previous exploratory MaAsLin2 analysis of the species level 16S rRNA amplicon sequencing data identified the same association of breast-feeding status on microbiota composition with a stronger effect with species level data (**Figure 3C; Supplementary File 6**). Species with relative abundance that was statistically significantly associated with breast feeding and recent diarrheal illness are shown as a heat map with positive and negative symbols indicating the direction of association (**Figure 2C** ). *Bifidobacterium longum* showed the strongest positive association with breast-feeding, with consistently higher median relative abundance in breastfed infants compared to those not breastfed (**Figure 3D** ) while *Klebsiella pneumoniae* showed the strongest negative association with breast feeding (**Figure 3E**). The species *Shigella flexneri* (**Figure 3F**) and *Clostridium perfringens* (**Figure 3G** ) were higher in samples taken after recent diarrheal illness. Breast feeding was the only variable associated with differences in microbial diversity at the species level of classification. The total number of bacterial species (**Figure 3H**) and the Shannon diversity was lower in breast fed infants than non-breast fed infants (**Figure 3I**). *Shigella flexneri* was detected in 49% of samples and *Clostridium perfringens* detected in 24% of samples indicating that these infants have a high prevalence of these potentially pathogenic genera in their gut microbiota (**Supplementary File 7**).

As described at the genus level, classification as stunted with a height-for-age z-score of less that 2 was not associated with gut microbiota species composition. Similarly, length-for-age z-score , underweight z-score, and weight-for-length z-score as continuous measures were also not associated with species composition. Hygiene measures of household toilet facilities and total number of people in the household were again not associated with species composition. Also noted was the lack of an association between the gut microbiota composition and recent antibiotic use (**Supplementary File 6**).

These species level results support the finding that breast feeding status was an major influence on the gut microbiota composition and associated with the abundance of *Bifidobacterium longum*. Recent diarrheal illness was associated with higher abundance bacterial species including *Shigella flexneri* and *Clostridium perfringens* that are known to be potential causes of diarrheal illness.

### Infants harbour bacterial strains that encode diverse human milk oligosaccharide and antimicrobial resistance genes

The frozen faecal samples that were not stored in DNA preservation solution were used to culture and isolate individual colonies for whole genome sequencing to gain further insight into the species of bacteria in the gut microbiota of these infants. Genomic analysis was carried out on the sequenced isolates to identify the species (and strains), and for the presence of virulence traits, antimicrobial resistance genes and HMO gene clusters. From the bacterial isolates genome sequenced 2 were identified as *Bacteroides*, 6 as *Bifidobacterium*, 1 as *Ruminococcus*, 3 as *Enterococcus*, 21 as *Clostridium*, 3 as *Clostridioides*, and 2 as *Paraclostridium* (**Figure 4A**).

**Figure 4.**
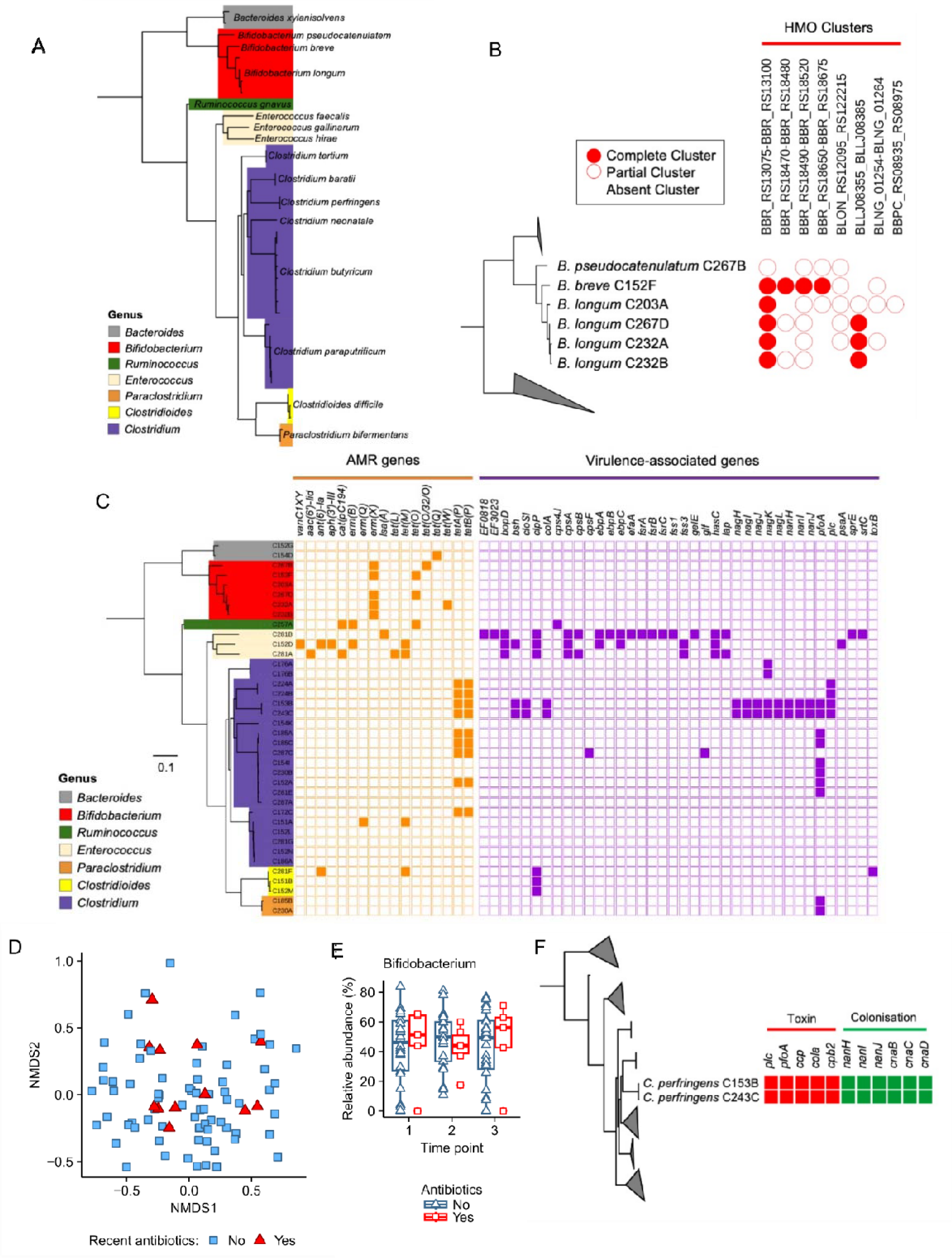
Genomic analysis of isolates for species determination, human milk oligosaccharide (HMO) gene clusters, antimicrobial resistance genes, and virulence genes. (A) Tree showing species and relatedness of whole genome-sequenced isolates; (B) Presence of HMO gene clusters in *Bifidobacterium* isolates; (C) Antimicrobial resistance genes and virulence associated genes identified in the isolate genomes; (D) NMDS plot showing all clustered faecal samples highlighted with recent antibiotic use; (E) *Bifidobacterium* genus abundance by antibiotic use; (F) Presence of toxin genes and colonisation factors in *Clostridium perfringens* isolates.

*Bifidobacterium* isolates comprised three species including *B. pseudocatenulatum*, *B. breve*, and *B. longum*. Both *B. breve* and *B. longum* isolates contained complete HMO gene clusters, with none in *B. pseudocatenulatum* (**Figure 4B**). This indicates the ability of the *B. breve* and *B. longum* present to digest the oligosaccharides present in human breast milk, with only *B. longum* showing higher relative abundance in breastfed infants compared to non-breast fed (**Figure 4C**). Comparing the four genome-sequenced bacterial isolates identified as *Bifidobacterium longum* using Average Nucleotide Identity with a 98% cut-off, three were identified as *B. longum* subsp. *longum* and one as *B. longum* subsp. *suis* (**Supplementary File 8; Supplementary Figure 1**).

Genomic analysis showed the presence of antimicrobial resistance genes across all strains isolated (**Figure 4D**). Of particular note was the presence of the antimicrobial resistance genes *erm*(X), *tet*(O), *tet*(O/23/O), *tet*(Q), and *tet*(W) in the genomes of isolated *Bifidobacterium* species (**Figure 4B** ), which provide resistance to erythromycin (*erm*) and tetracycline (*tet*) respectively. The identification of antimicrobial resistance genes in the *Bifidobacterium* strains present may link with the lack of any association between gut microbiota composition and those infants receiving antibiotics, as shown here with NMDS plots showing all samples combined (**Figure 4E** ). The relative abundance of

*Bifidobacterium* was unchanged in samples from infants receiving antibiotics (**Figure 4F** ). Amongst this group of infants the most commonly received antibiotic was amoxycillin, with individual instances of cefixime, and cefadroxile also received (**Supplementary File 2**).

Two *Clostridium perfringens* strains were isolated and genome-sequenced. Both contained five toxin producing genes and six colonisation factor genes that relate to the ability to cause infection and disease in the gut (**Figure 4G**). This confirms the presence of *C. perfringens* that was identified using 16S rRNA amplicon sequencing and the presence of toxin and colonisation factor genes supports a potential role in its association with cases of infant diarrhoea.

### Faecal cytokines profiles are associated with breastfeeding status

The early life gut microbiota is known to play a key role in shaping gut and mucosal immune responses. A multi-plex assay including a panel of thirty cytokines was used to determine concentrations in the infant faecal samples (**Supplementary File 9**).

A linear mixed model incorporating infant metadata variables as a fixed effect and the infant as a random effect was used as an exploratory analysis of associations between each cytokine and metadata variables in all sample across the three time points (**Supplementary File 10** ). The most robust and consistent association was identified between two cytokines and breast feeding (**Figure 5A-C**). The cytokines Interleukin-1-alpha (IL-1α) (**Figure 5A** ) and Vascular endothelial growth factor (VEGF) (**Figure 5B** ) were at significantly higher concentration in breast-fed infants. Both cytokines showed substantial variation in concentration across time points within samples from the same infant. IL-1α (**Figure 5A** ) and VEGF (**Figure 5B** ) were also the two cytokines with the highest concentrations in the faecal samples across all infants.

**Figure 5.**
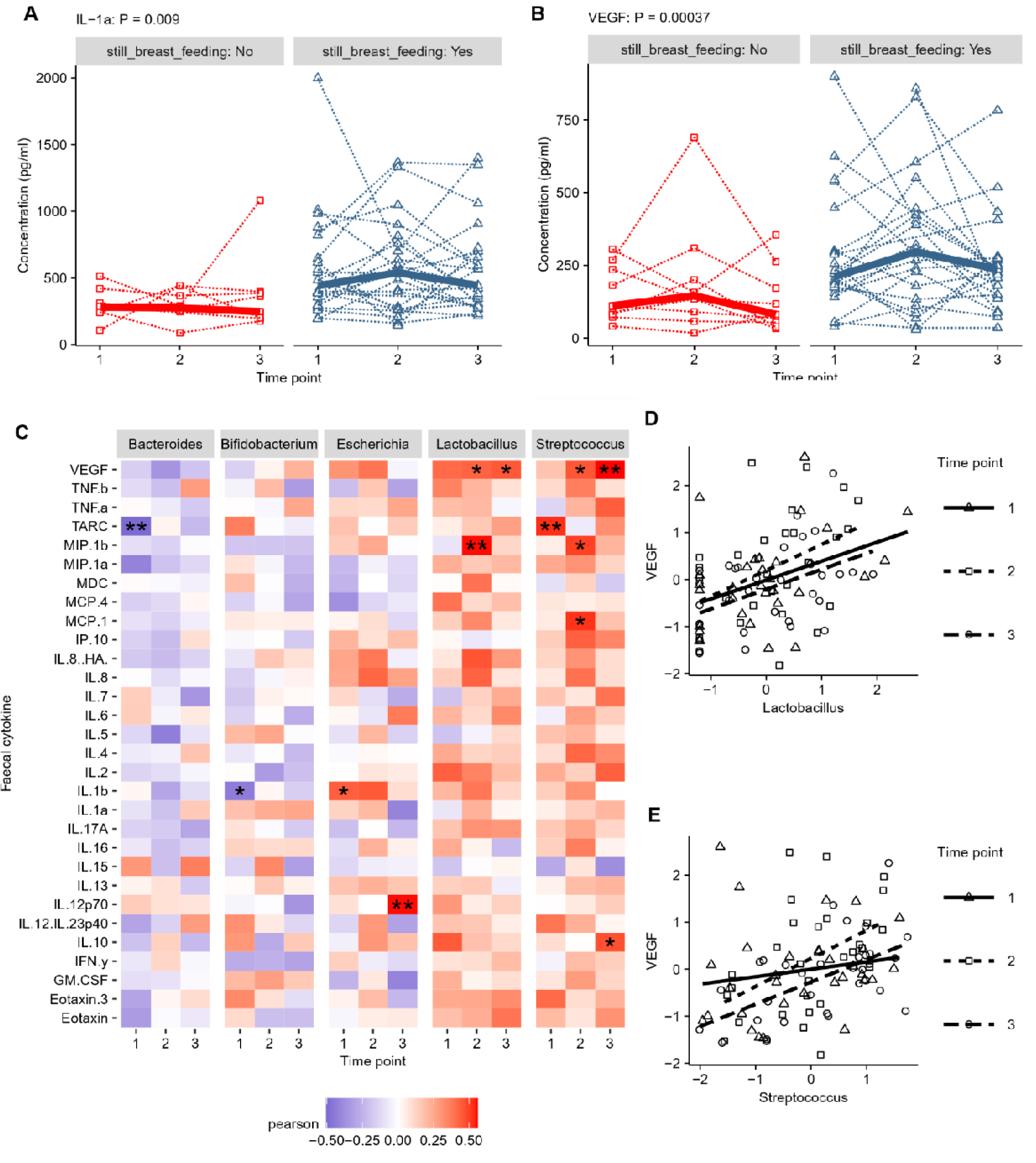
Infant faecal cytokines concentration and associations with breast feeding status. (A) Individual infant IL-1α concentration by breastfeeding status across time points; (B) Individual infant VEGF concentration by breastfeeding status across time points; (C) Pearson correlations between faecal cytokine concentrations and five highest abundance genus with variables in both datasets individually normalised using the bestNormalize package; (D) Scatter plot of VEGF and *Lactobacillus* correlations labelled for each time point; (E) Scatter plot of VEGF vs *Streptococcus* correlations labelled for each time point.

To identify whether differences in major faecal cytokines by breastfeeding status were linked to differences seen in the microbiota composition a correlation analysis was carried out between faecal cytokine concentrations and the relative abundance of the 5 most abundant genus (**Figure 5C**) and species (**Supplementary Figure 2** ) in the microbiota. Weakly consistent positive associations were seen between VEGF and *Lactobacillus* (**Figure 5D** ) and *Streptococcus* (**Figure 5E** ) across the three time points, significant at two time points after correction for multiple testing. There were no consistent correlations between the concentration of other cytokines and relative abundance of individual bacterial genus or species, including *Bifidobacterium*.

## Discussion

This study represents the first time that the infant gut microbiota composition of a cohort of rural Cambodian infants has been analysed in detail. We show that shortage of samples under rural conditions can radically reduce the abundance of certain genus within the samples. Infants had a gut microbiota dominated by the genus *Bifidobacterium,* shaped to a large degree by breastfeeding, which also impacts faecal cytokine profiles. Previous gut microbiota research in Cambodia has included investigation into antimicrobial resistance genes in Enterobacterales (Auguet et al., 2021) the epidemiology of antibiotic resistance in *Escherichia coli* and *Klebsiella pneumoniae* (van Aartsen et al., 2019), and effects of iron supplementation on individual genus abundance in women (Finlayson-Trick et al., 2023). This present study goes further and examines in-depth genus and selected species and strain composition in Cambodian infants for the first time.

Our study results represent an unusual example of different sample storage conditions as these samples were stored without immediate freezing. Overall the effect of storage conditions on the composition of the microbiota was small but a higher relative abundance of *Bacteroides* in particular was seen in samples stored in DNA preservation solution and a lower abundance of common species including *Bifidobacterium* due to the proportional nature of the microbiota data. *Bacteroides* represents an important genus in the infant gut microbiota, transferred from the mother during delivery and has been found to be higher in vaginally delivered infants (Matharu et al., 2022) with some *Bacteroides* species having the ability to consume components of breast milk (Kijner et al., 2022). The relatively high proportion of *Bacteroides* in these infants and their almost disappearance from samples without DNA preservation solution shows the importance of a storage method that preserves this significant component of the infant microbiota.

Previous research has investigated the effects of storage conditions on microbiota analysis and reported alterations in *Bacteroides* abundance, although this has focused on storage conditions while frozen or number of freeze thaw cycles (Pérez-Burillo et al., 2021; Poulsen et al., 2021). Room temperature storage of faecal samples for up to 72 hours has been reported to reduce genus including *Bacteroides* in adult faecal samples over time (Roesch et al., 2009). While defrosting frozen samples for three hours before analysis reduced *Bacteroides* abundance by half (Cardona et al., 2012). The duration of cool but unfrozen temperatures that samples in the present study were exposed to appear to have resulted in the particular low of *Bacteroides*. However, the unpreserved samples were also important for culturing and isolating live bacteria for genomic sequencing, highlighting the importance of having samples stored under both conditions.

*Bifidobacterium* was present at lower abundance in non-breast fed infants and not breast feeding was associated with higher abundance of potentially pathogenic bacterial genus like *Klebsiella* and *Clostridium*. Although the numbers of infants not breast fed was too low to draw strong conclusions, previous research has indicated that increased *Bifidobacterium* in the preterm infant gut can help reduce the proportions of potential pathogens (Alcon-Giner et al., 2020). This is linked to the ability of *Bifidobacterium* to metabolise HMOs in breast milk and reduce gut pH to make it a more inhospitable environment for overgrowth of opportunistic pathogens. *Bifidobacterium longum* was the dominant species in this cohort of infants and *B. longum* was the only *Bifidobacterium* species with higher abundance in breast fed infants. From the four *B. longum* isolates genome-sequenced, all contained one or two HMO gene clusters indicating that they were potentially capable of metabolising the sugars present in breast milk. Interestingly, three of the *B. longum* isolates were confirmed to be *B. longum* subsp. *longum*, which do not typically dominate the gut microbiota of infants in developed western countries nor other LMICs with high rates of breast feeding. It was recently reported that *B. longum* subsp. *infantis* was the dominant *Bifidobacterium* in 1 to 2 month old infants from the Gambia and Bangladesh with *B. longum* subsp. *longum* only forming a minor proportion of the microbiota (Taft et al., 2022). The isolation of *B. longum* subsp. *longum* from Cambodian samples may be due to sampling at an older age of around 7 months, with recent research from Bangladesh indicating that a transition occurs from *B. longum* subsp. *infantis* to *B. longum* subsp. *longum* with increasing infant age (Vatanen et al., 2022). One *B. longum* isolate was identified as most closely matching *B. longum* subsp. *suis*, a subspecies originally isolated from pigs (Matteuzzi et al., 1971) and found to be a dominant member of the piglet gut microbiota (Zani et al., 1974). *Bifidobacterium* isolates corresponding to *B. longum* subsp. *suis* were recently identified in Bangladeshi infants (Vatanen et al., 2022), which suggests that this may be an under-recognised member of the infant gut microbiota. From only a single isolate we cannot determine the proportion of *B. longum* subsp. *suis* within Cambodian samples, and this could represent the transient colonisation of this bacterium from close proximity to pigs or that this is a resident of the gut microbiota in this region. The presence of an HMO gene cluster capable of metabolising breast milk sugars indicates that this *B. longum* subsp. *suis* isolate may be capable of metabolising breast milk HMOs. The dominant abundance of the species *B. longum* in breastfed infants underline the central effect of diet on the gut microbiota composition of these infants, overwhelming any other environmental factor in this small cohort.

Previous research has identified associations between restricted infant growth and stunting due to malnutrition and differences in the infant gut microbiota (Monira et al., 2011; Surono et al., 2021). We did not find any associations between the gut microbiota and infants growth. This was potentially due to the small cohort of infants involved with only three out of the thirty infants included showing stunted growth and the relatively young age of the infants starting at an average of 7 months of age. Previous research has looked at older children (Monira et al., 2011) or older infants with a higher proportion of the infants suffering from stunting (Surono et al., 2021). This indicates that in this small cohort of infants it is the presence or absence of breast milk having the largest effect, potentially overwhelming the ability to detect other environmental influences.

In Cambodia, and many other LMICs, infants can be exposed to pathogens through provision of contaminated drinking water in the first months of life (Poirot et al., 2018). Whilst we show that there are high proportions of beneficial bacteria in most samples, pathogenic taxa including *Shigella*, *Escherichia*, and *Clostridium* genus were also detected in most infants. Although *Shigella flexneri* and *Clostridium perfringens* abundance in the gut was associated with recent diarrhoea these bacterial species were also detectable in a large minority of infant samples indicating environmental exposure to these organisms and potential asymptomatic carriage.

Antibiotics can significantly alter the profile of the early gut colonization (Alcon-Giner, et al., 2019) resulting in reduced diversity (Gallego et al. 2019) and susceptibility to other infections (Porras and Brita 2019). Notably, in Cambodia there is widespread, unregulated and inappropriate antibiotic use, and these selection pressures may further facilitate the dissemination of AMR genes (van Aartsen et al., 2019). An estimate of one-third of the Cambodian population is < 18 years old, and this group may be a significant reservoir for the spread of antimicrobial resistant organisms (van Aartsen et al., 2019). Indeed, we observed multiple antibiotic resistance genes in potentially pathogenic species, including those associated with diarrhoeal episodes. We also observed presence of antimicrobial resistance genes in *Bifidobacterium*. Although this may seem unexpected, as *Bifidobacterium* is very rarely reported to be antibiotic resistant, a recent report from the neighbouring country of Vietnam indicates that *Bifidobacterium* isolated from healthy infants harbour antibiotic resistance genes (Chung The et al., 2021). We also found that Cambodian *Bifidobacterium* isolates contained resistance genes to the antibiotics erythromycin and tetracycline. This may link to the fact that this cohort of infants has previously been reported to be exposed to frequent and unregulated use of antibiotics (Miyazaki et al., 2020), thus potentially driving AMR dynamics in beneficial members of the infant gut microbiota. However, although amoxicillin was the most commonly used antibiotic in this cohort, we did not observe any beta-lactam resistance genes in the isolates. Previous studies have indicated that while it appears there is high and unregulated use of antibiotics within the Cambodian population – they may be ineffective due to inappropriate storage thus reducing their effectiveness. However, the widespread presence of resistance genes in other isolates may point towards active selection through exposure to antibiotics. The apparent maintenance of *Bifidobacterium* abundance despite antibiotic treatment may point to the presence of other novel resistance genes in these *Bifidobacterium*, which if identified may have future potential for combining effective probiotic therapy with antibiotic treatment.

One genus notable by its absence from this cohort of Cambodian infants is the genus *Segatella*. Previously found to be a dominant member of the gut microbiota in rural West Africa while rare in Italian children (De Filippo et al., 2010). More recently this high prevalence of *Segatella* in children in low income countries has been confirmed in The Gambia, Kenya, Mali, and Bangladesh (Pop et al., 2014). Whilst *Segatella* abundance was lower in children from Bangladesh it was still detectable in children of a similar age to this cohort of Cambodian infants (Pop et al., 2014). In contrast Cambodian infants contained *Bacteroides*, a genus previously associated with the gut microbiota of European and US inhabitants (De Filippo et al., 2010). What dietary differences may account for this difference cannot be determined in this study, however, it does show that further investigation is needed into underrepresented populations. Whether the presence of *Bacteroides* and absence of *Segatella* persists into adulthood remains unknown as composition of the adult gut microbiota in this region remains still to be investigated.

Consistent differences in the faecal concentrations of the cytokines IL-1α and VEGF point to potential effects of breastfeeding on the infant gut mucosal immune system. VEGF has been reported to be secreted by the mammary glands, and breast milk contains high concentrations of free VEGF that can bind to receptors on the surface of cells in the infant intestine (Kobata et al., 2008; Siafakas et al., 1999; Vuorela et al., 2000). Reduced VEGF in preterm infants is suspected to be involved in necrotising enterocolitis which is associated with overgrowth of opportunistic pathogens including *C. perfringens* (Kiu et al., 2023; Sabnis et al., 2015). While the role of VEGF originating from breast milk in term infants remains unexplored this growth factors could exert growth-promoting and protective effects on the infant gut (Kobata et al., 2008). While the role of IL-1α in the infant gut has not been explored, it has also been reported to be present in breast milk (Zanardo et al., 2005) however, whether this has any effects the infant gut is currently unknown. Although we are unable to demonstrate a breast milk origin from these results, the presence of higher concentrations of these three faecal cytokines in breast fed infants does suggest that they may be derived from breast milk, with a potential to influence infant development that warrants further investigation.

While there was little consistent association across time points between the gut microbiota at the genus level and faecal cytokines, a weak positive correlation was noted between *Lactobacillus* and *Streptococcus* and the faecal cytokine VEGF. As *Streptococcus* abundance was higher in breastfed infants this correlation is potentially linked to breastfeeding. As both microbiota composition and faecal cytokines is variable with time it may not be possible to identify links between factors in a limited sample size of infants. Interestingly, despite being the genus most differentiated by breast feeding status there no consistent correlations between *Bifidobacterium* IL-1α or VEGF. Indeed, many species and strains from this genus are known to positively influence the gut and systemic immune system – including *B. longum,* which is proposed to be in part mediated by metabolites produced as a result of HMO degradation e.g. acetate (Henrick et al., 2021; Laursen et al., 2021). However, further studies including larger number of infants would need to be performed to explore microbial-immune interactions in more detail. In addition, the use of the frozen samples for faecal cytokine analysis may be limited as it is unknown how such storage conditions may impact the degradation of the cytokines tested.

We show that this cohort of infants from rural Cambodia have a *Bifidobacterium* dominated gut microbiota (including atypical species) that is primarily shaped by breastfeeding. This highlights the influence of breast feeding in supporting a beneficial infant microbiota in rural conditions. As *Bacteroides* represents a common and important genus in the human gut the effects of sample storage are an important consideration when immediate freezing of samples is not possible. The lack of impact of antibiotic use may indicates either widespread antibiotic resistance or the use of ineffective antibiotics. The finding that faecal cytokines are shaped by breast feeding and could potentially originate from breast milk is a finding with unknown but potentially interesting implications. This study adds significantly to the available data on the composition of infant gut microbiota communities in a previously neglected and difficult to sample geographical area of the world. Moreover, given Cambodian profiles are somewhat different from other infant cohorts, including LMICs, studies such as these shows the importance of looking at infants from underrepresented areas for defining what can constitute a normal infant microbiota.

## Materials & Methods

### Study design & ethics

Samples were collected from a sub-group of infants enrolled in NHAM cohort based in villages along the Mekong river in rural Kampong Cham province in Cambodia. This registered all infants born between April 2016 and March 2019 with the objective of investigating causes of stunting. Full ethical approvals including National Ethical Committee for Health Research in the Ministry of Health, Cambodia (016) and the Ethical Committee of the National Centre for Global Health and Medicine in Japan (NCGM-G-001870-01), and methods for the collection of detailed meta-data for infants included in this sub-study are as previously described (Miyazaki et al., 2021, 2020). Weight and length were measured to the nearest 0.1kg and 1 mm by trained field workers at each home visit using a weighing scale (Seca 877, Seca, Hamburg, Germany) and a length board (Seca 417, Seca).

### Sample collection

Infant faecal samples were analysed from a subset of 32 infants out of 47 enrolled in a detailed 3- month longitudinal follow-up study (Miyazaki et al., 2021, 2020) collected between March and July in 2017. Mothers/primary care-givers and their infants were visited by study field workers every 14 days. Three samples were analysed from each infant for a total of 96 individual samples. The trained field worker gave instructions on how to collect and equipped the caregivers with the material for the child stool collection prior to routine home visits. Once stool sample collection by mothers or caregivers was completed, the sample was kept in relatively cooler place in the house or in cooler bags with ice until the field worker visited to collect them. The majority of the samples were collected by the field worker within 10 hours of the time of child defecation reported by mothers (maximum: 47 hours and minimum: 15 min among all sample). Each sample collected was divided in two and half stored in DNAShield and stored unaltered at ambient temperatures until delivered to laboratory. The other half of the sample was stored in iceboxes with refrigerant and transported to a -20° freezer on the same day and once delivered was stored at -80°. The first sample was collected after an initial visit where cross-sectional metadata was collected. Subsequent samples were collected at approximately two week intervals along with further individual visit metadata. Underweight z-scores (weight-for-age), weight-for-length z-scores, length-for-age z-score were calculated previously (Miyazaki et al., 2021). A hand washing score was generated as a composite score combining the frequency and consistency of handwashing with soap. Improved toilet facility was defined as having access to a toilet flushing to a septic tank or pit latrine compared to unimproved having only access to a pit latrine or no toilet facility. Household toilet facility was defined as having a toilet within the home flushing to a septic tank.

### DNA extraction of preterm stool samples

FastDNA Spin Kit for Soil (MP) was used to extract DNA from preterm faeces following manufacturer instructions, with extended 3 min bead-beating. DNA concentration and quality were quantified using a Qubit® 2.0 fluorometer (Invitrogen).

### 16S rRNA amplicon sequencing of faecal samples

16S rRNA region (V1-V2) primers were used for library construction as described previously (Alcon-Giner et al., 2020). This set of primers allowed the amplification of one 16S rRNA gene sequencing library containing 96 different samples. PCR conditions used were: cycle of 94°C 3 min and 25 cycles of 94°C for 45 s, 55°C for 15 s and 72°C for 30 s. Sequencing of the 16S rRNA gene libraries was performed using Illumina MiSeq platform with 300 bp paired end reads.

### 16S rRNA amplicon data analysis

The raw sequence reads obtained were subjected to quality filtering using trim_galore v0.4.3. The resulting filtered sequence reads were then mapped against the 16S rRNA SILVA database (SILVA_132_SSURef_tax_silva) using BLASTn v2.2.25+ with a maximum e-value threshold of 10e-3. This mapping process was performed separately for both pairs of sequences (Quast et al., 2013). After completing the BLASTn alignments, the output files were annotated using the paired-end protocol of MEGAN6, employing the default Lowest Common Ancestor (LCA) parameters (Huson et al., 2016; Huson and Mitra, 2012).

Statistical analysis was conducted using R version 4.2.2 (2022-10-31 ucrt) and RStudio (2023.06.0+421). To determine the number of reads required to adequately represent the microbiota in each sample, rarefaction curves were generated using the vegan package in R. For the analysis at the genus level, the sample with the smallest number of reads contained 37,306 classified reads, which was deemed an acceptable minimum. For the species level analysis, the sample with the second lowest number of reads contained 9,317 reads, which was considered an acceptable minimum. The 16S rRNA gene sequence data were then normalized by subsampling using the rarefy function in the phyloseq package, resulting in an even depth of 37,306 reads for the genus data and 9,317 reads for the species data. One sample was excluded from the species data analysis as it only contained 3,965 reads.

To examine differences in the relative abundance of genera among the 69 infant fecal samples, which had 16S rRNA sequence data obtained from both samples stored in DNAShield solution and Frozen, a Wilcoxon rank sum test was performed in R. This test was chosen due to the non-parametric nature of the majority of the genus abundance data. To account for multiple testing, a Benjamini & Hochberg false discovery rate (FDR) method was applied, and a p-value of less than 0.05 was considered statistically significant for adjusted p-values.

The overall composition of the 16S rRNA gene sequence data was analyzed using NMDS (Non-metric multidimensional scaling) plots. These plots were generated in R Studio using the vegan package version 2.6-4, employing a Bray-Curtis dissimilarity calculation. To determine the statistical significance of differences in microbial community structure based on infant metadata variables, permutational MANOVA was conducted using the Adonis function from the vegan R package version 2.6-4.

Microbial diversity at both the genus and species taxa levels was calculated using the vegan package version 2.6-4. The number of genera or species was determined for each sample, and the Shannon diversity index was calculated accordingly. Linear mixed-effect models, utilizing the “lmer” function in R, were employed to test the associations between the relative abundance of genera or species and metadata variables. The number of genera or species, as well as the Shannon diversity index, were assessed against each individual metadata variable, with Time Point included as a fixed effect and Patient ID included as a random effect in the model.

To test for differential abundance of genera and species in relation to available background metadata variables for the infant samples, the Maaslin2 package (Microbiome Multivariable Associations with Linear Models) (Mallick et al., 2021), was utilized. The Maaslin2 analysis was performed with default parameters, including Benjamini-Hochberg FDR multiple testing correction and a significance threshold of 0.25. No minimum prevalence or abundance cut-offs were applied, and the data input was previously normalized. Each metadata variable was individually tested, incorporating Time Point and Patient ID as random effect grouping variables in a mixed-effect model. Positive associations were identified between Breast Feeding and Diarrheal Illness. These two variables were subsequently tested together as fixed effects, with Time Point and Patient ID included as random effects in the final analysis.

### Anaerobic bacterial culturing and isolation from infant faecal samples

Anaerobic bacterial culturing and isolation was performed on seventeen Cambodian infant faecal samples (C151, C152, C153, C154, C172, C176, C185, C186, C203, C224, C230, C232, C243, C257, C267, C281, and C287) following (Browne, 2018; Browne et al., 2016; Forster et al., 2019). Sample preparation was performed under anaerobic conditions. YCFA media and PBS was pre-reduced in an anaerobic cabinet at 37 °C for a minimum of 12hs before culturing. Two samples, weighing 100mg each, were prepared from each faecal sample. The first sample was homogenised in reduced PBS, serially diluted, and plated directly onto the YCFA agar. The second sample was ethanol treated using 70% ethanol under ambient aerobic conditions for 1h to kill vegetative cells; the sample was then washed three times with reduced PBS to remove the ethanol, homogenised in the reduced PBS, serially diluted and plated onto YCFA + Sodium taurocholate agar to stimulate spore germination. All agar plates were placed in the anaerobic cabinet at 37°C for 48hs. Single colonies were picked and streaked onto pre-reduced YCFA media to isolate and purify the colonies; YCFA media was pre-reduced in the anaerobic cabinet at 37 °C for a minimum of 12hs before isolating colonies. Colonies from the non-ethanol treated samples were streaked onto the reduced YCFA agar plates, and colonies from the ethanol treated samples were streaked onto the reduced YCFA + Sodium taurocholate agar plates. Plates were then left to incubate in the anaerobic cabinet at 37°C for 48-72hs. This was carried out a second time to make sure isolates were pure. Falcon tubes containing reduced YCFA broth was inoculated with a single colony from the YCFA and YCFA + Sodium taurocholate agar plates, and left to incubate in the anaerobic cabinet at 37°C for 48hs, with bacterial stocks prepared in 1.5ml cryovials with 30% glycerol YCFA broth, and stored at -80°C. Additionally, two infant faecal samples C203 and C232 were processed for targeted *Bifidobacterium* isolation. Briefly, serially diluted aliquots of faecal homogenates (10^-1^ – 10^-6^ in PBS) were plated onto MRS agar supplemented with 50mg/l mupirocin and 0.5g/l L-cysteine HCL and incubated in an anaerobic cabinet at 37 °C for 48h. Colonies with desired morphology were randomly selected and re-streaked on MRS agar to purity. Pure cultures were prepared in 1.5 ml cryovials with 30% glycerol Reinforced Clostridial Medium (RCM) and stored at –80 °C.

### Whole Genome Sequencing (WGS)

DNA extractions were performed using the MPbio Fast DNA Spin Kit for soil and bead beating (but with an extended 3 min bead beating time). DNA quantification was carried out using the Qubit 2.0 fluorometer. DNA from pure bacterial cultures was sequenced at Wellcome Trust Sanger Institute using 96-plex Illumina HiSeq 2500 platform to generate 125 bp paired end reads as described previously (Harris et al., 2010).

### Genomic analysis

Paired-end raw sequence reads (fastq) were quality-filtered (-q 20) via fastp v0.20.0 (Chen et al., 2018) prior to de novo genome assembly using SPAdes v3.14.1 (Bankevich et al., 2012) at default parameters. Next, contigs smaller than 500bp were discarded in each genome assembly. All genome assemblies were subjected to contamination check via CheckM v1.1.3 (Parks et al., 2015), contaminated (contamination >5%) and/or incomplete (completeness <90%) genome assemblies were excluded from further analyses (n=5). All genomes were identified taxonomically using gtdb-tk v1.5.1 (Chaumeil et al., 2020), using ANI cut-off 95% against type strains of each species.

These high-quality genome assemblies (n=71) were then dereplicated at ANI 99.9% (cut-off for strain level) using dRep v3.2.2 (Olm et al., 2017) prior to generation of a distance tree using Mashtree v1.2.0 (Katz et al., 2019) on default parameters. Tree was annotated in iTOL v6.5.8 (Letunic and Bork, 2019).

These genomes were also subjected to sequence search analysis via ABRicate v1.0.1 (Seemann, 2023) based on ResFinder v4.0 database (Bortolaia et al., 2020) (--minid=95 and --mincov=90) to determine antimicrobial resistance (AMR) profiles, and VFDB database (Liu et al., 2019) (--minid=90 and –mincov=90) to determine virulence profiles. Furthermore, HMO clusters were detected using an in-house HMO sequence database via ARBricate at –minid=70 (Lawson et al., 2020). In addition, *C. perfringens* virulence profiles were predicted via database TOXIper v1.1 using ABRicate at – minid=90 (Kiu, 2023).

For *Bifidobacterium longum* subspecies analysis, *B. longum* strains were compared with four subspecies type strains namely subspecies *infantis*, subspecies *longum*, subspecies *suillum* and subspecies *suis*, to generate a core gene alignment via Panaroo v1.2.8 (Tonkin-Hill et al., 2020), resulting in 1,212 core genes, which core-gene alignment constructed by MAFFT (Katoh and Standley, 2013) were SNP-extracted (via SNP-sites v2.3.3)(Page et al., 2016) and subsequently used to generate a phylogenetic tree via IQ-TREE v2.0.5 (Minh et al., 2020, p. 2) with ultra-fast bootstrap replicates -B 1000 and automatic evolutionary model selection option -m TEST (best fit model was determined to be GTR+F+ASC+G4). Subspecies identification was performed using fastANI v1.3 (Jain et al., 2018) at cut-off 98%.

### Faecal cytokine measurement

Faecal samples were homogenized with PBS using a FastPrep® Bead Beater (4.0m/s, 3min), centrifuged (14,000rpm, 15min) and 25μl of supernatant was used for the assay. Samples were analysed using MULTI-SPOT™ plates, MESO Quickplex SQ120 and discovery workbench software according to the manufacturer’s protocol. Pre-coated immunoassays V-PLEX Proinflammatory Panel 1 (human) and V-PLEX Cytokine Panel 1 (human) were used to detect a set of 20 different cytokines: IFNγ, IL-1β, IL-2, IL-4, IL-6, IL-8, IL-10, IL-12p70, IL-13, TNFα, GMCSF, IL-1α, IL-5, 273 IL-7, IL-12p40, IL-15, IL-16, IL-17A, TNFβ, and VEGF-A.

### Data and Code Availability

Bacterial pure genome assemblies are now deposited at NCBI GenBank under project accession PRJNA958097 while 16S rRNA amplicon sequencing raw reads are deposited in NCBI SRA under project PRJNA958098.

### Statistical analysis

Associations between faecal cytokine concentrations and metadata variables were tested using the MaAsLin2 package (Mallick et al., 2021), a differences in concentration against background metadata variables available for the infant samples. For faecal cytokine analysis MaAsLin2 was run with default parameters including multiple testing correction using Benjamini-Hochberg FDR with a significance threshold of 0.25, except: no minimum prevalence or abundance cut-offs were applied, normalised methods used wans “CLR”, and the transformation “LOG”. Each metadata variable was tested individually with Time Point and with Patient ID included as a Random Effect grouping variable to run a mixed effect model. Positive associations were identified with Breast Feeding and Diarrheal Illness and these two variables where tested together as Fixed Effects with Time Point and Patient ID included as a Random Effect in the final analysis.

Correlations between faecal cytokine concentration and the relative abundance of genus and species was tested using a Pearson correlation with correction for multiple testing using Benjamini & Hochberg FDR correcting between both cytokines and time point. Both cytokine concentration and genus/species abundance were first normalised using the bestNormalize package in R.

## Author contributions

Conceptualisation, L.J.H and S.E.C.; Methodology, M.J.D, R.K., I.R.S, A.M., HA-P., R.T., S.C., S.P., M.K., M.M., A.I., and B.T; Software, M.J.D., R.K., S.C., and M.K.; Validation, M.J.D, K.S., S.E.C., and L.J.H.; Formal analysis, M.J.D, R.K., A.M, S.E.C., and L.J.H.; Investigation, M.J.D, R.K., I.R.S, A.M., HA-P., R.T., S.C., S.P., M.K., M.M., A.I., B.T., S.E.C., and L.J.H.; Resources, M.J.D., A.M., R.T., M.M., A.I., B.T., S.E.C., and L.J.H; Data Curation, M.J.D., R.K., A.M., S.E.C.; Writing - Original Draft, M.J.D. and L.J.H.; Writing - Review and Editing, M.J.D, R.K., I.R.S, A.M., HA-P., R.T., S.C., S.P., M.K., M.M., A.I., B.T., S.E.C, and L.J.H; Visualisation, M.J.D and R.K.; Supervision, S.E.C,. and L.J.H.; Project Administration, M.J.D., S.E.C, and L.J.H.; Funding Acquisition, S.E.C, and L.J.H.

## Acknowledgments

We thank the families for participating in the study.

## Supplementary Files

Supplementary file 1. Genus differences between DNAShield and Frozen storage conditions.

Supplementary File 2. Metadata and Genus Sequence Data

Supplementary file 3. Genus data Maaslin2 analysis results.

Supplementary file 4. Genus prevalence in samples.

Supplementary file 5. Combined sample metadata and species sequencing data for all samples.

Supplementary file 6. Species data Maaslin2 analysis results.

Supplementary file 7. Species prevalence in samples.

Supplementary File 8. Pairwise Average Nucleotide Identity (ANI)

Supplementary file 9. Metadata and Faecal Cytokine Concentrations.

Supplementary file 10. MaAsLin2 Cytokine Results

## Supplementary Figures

**Supplementary Figure 1.**
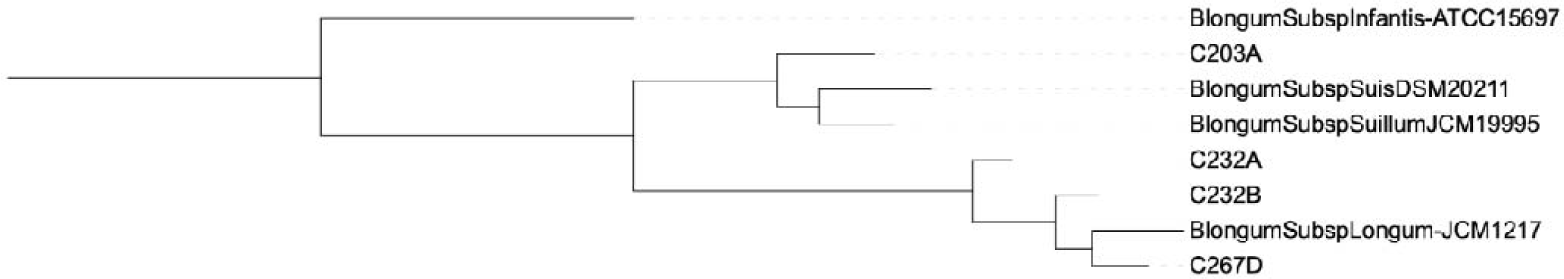
The subspecies of Bifidobacterium longum isolates determined using average nucleotide identity with a 98% cut-off.

**Supplementary Figure 2.**
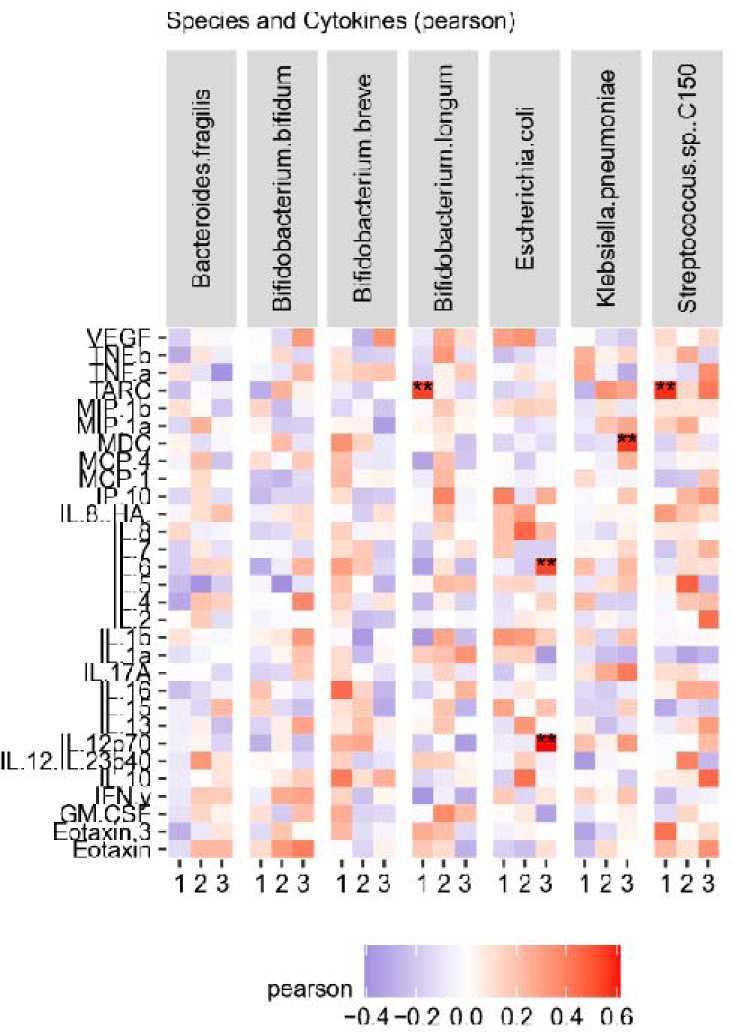
Correlations between cytokines and Genus or Species relative abundance.

